# Emergent directional persistence in fibrous granular scaffolds guides myotube organization

**DOI:** 10.64898/2026.05.01.721636

**Authors:** James L. Gentry, Steven R. Caliari

## Abstract

Granular scaffolds have emerged as promising platforms for tissue regeneration, offering injectability and cell-scale porosity that support robust cell infiltration and tissue formation. However, the isotropic pore structure of spherical building blocks does not provide the directional cues needed to guide organized tissue formation. Addressing this requires asking not just whether granular scaffolds can be made anisotropic, but whether directional cues persist across the pore network at scales relevant to cell behavior. Using high aspect ratio GelMA hydrogel fibers as building blocks, we demonstrate that spherical granular materials lose orientational coherence at the cellular scale, confirming that isotropic building blocks are fundamentally incapable of providing structural guidance beyond individual pore neighborhoods. In contrast, fibrous building blocks extend persistence into the multicellular range, occupying an intermediate architectural regime exhibiting locally coherent but globally variable organization, rather than simple isotropic or uniaxial alignment, that has previously been inaccessible to granular scaffold design. We show this regime is functionally meaningful: myotubes undergo contact guidance through locally persistent but globally variable pore structure, and greater persistence is associated with increased myotube elongation and multinucleation in primary human muscle progenitor cells. Together these results expand the design space for granular scaffolds beyond pore size and porosity, and establish persistence as a variable linking granular scaffold architecture to organized tissue formation.

## Introduction

Granular scaffolds, constructs assembled from packed microgel building blocks, have emerged as a versatile platform for tissue engineering, combining the injectability and modularity of particle-based systems with the cell-scale porosity that supports robust cell infiltration and tissue formation^[5]^. Their pore architecture, formed by the interstitial space between packed microgels, is central to their function: pore size and porosity govern cell access and nutrient transport^[6]^, and these parameters are highly tunable through particle size, particle shape, and packing density^[7–10]^. Granular scaffolds have demonstrated strong regenerative outcomes across a range of tissues as a result of this inherent pore structure^[1,11,12]^. However, pore size and porosity describe only the dimensions of available space, and cannot describe any coordinated spatial organization of the porous microstructure.

The architectural properties that govern multicellular organization in scaffold systems extend beyond the geometry of individual pores to the collective organization of the pore network, or how pore orientation is maintained and coordinated across space. This emergent structure has proven critical for guiding cell organization in aligned electrospun fiber mats^[13–16]^, freeze-dried scaffolds with directional pore channels^[17–22]^, and micropatterned substrates^[23–26]^, all of which demonstrate that sustained directional cues guide cell organization, alignment, and migration over extended distances. Analytical tools have been developed to quantify this organization: local thickness to characterize pore dimensions^[27,28]^, structure tensor analysis to measure local pore orientation^[29,18]^, and nematic order metrics to quantify how consistently that orientation is maintained across space^[30–32]^. Granular scaffolds have been largely excluded from this design space and analytical tradition because isotropic spherical building blocks produce pore geometries in which orientation varies randomly from pore to pore, precluding sustained directional organization^[1,2]^. The absence of any preferred orientation implies that directional cues decorrelate rapidly over short distances, limiting the scaffold’s capacity to organize cells beyond the immediate pore neighborhood^[3]^.

This limitation has functional consequences in tissues that require spatially organized cellular architecture. Many native tissues exhibit locally coherent but globally variable organization rather than simple isotropic or uniaxial alignment^[4]^, suggesting that the relevant design target is persistent local organization. In skeletal muscle specifically, myofibers exhibit locally organized but spatially varied orientations that collectively enable efficient force generation and multiplanar movement^[33,34]^, and recapitulating this architecture during regeneration requires persistent directional cues at the supracellular scale^[35,36]^. Granular scaffolds have shown remarkable regenerative capacity in traumatic muscle injury models, supporting robust tissue formation and favorable immune modulation^[10,11,37]^. However, the only study to measure functional recovery following granular scaffold treatment found no improvement over untreated controls, attributed to the isotropic pore structure of spherical building blocks which produced tortuous, disorganized myofibers incapable of coordinated force generation^[11]^. These results suggest that robust tissue regeneration alone is insufficient without architectural cues to guide organization at supracellular length scales.

Introducing anisotropic building blocks is a natural strategy for imparting directional pore structure to granular scaffolds^[3]^. One line of work has focused on porosity gains, demonstrating that high aspect ratio particles pack into scaffolds with substantially higher void fractions than spherical building blocks, improving cell infiltration and scaffold permeability^[9,38]^. A separate study investigated pore anisotropy, showing that rod-shaped particles produce pores with higher aspect ratios than spherical ones, with corresponding improvements in cell invasion attributed to directional contact guidance cues^[7]^. However, pore aspect ratio describes individual pore geometry in isolation and does not address whether orientation is consistent between neighboring pores. Others have demonstrated aligned granular-like materials, albeit using fabrication approaches that prioritize permanent uniaxial alignment at the expense of the locally persistent but globally variable architectures characteristic of most native tissues^[39–42]^. While these studies establish that building block geometry influences local pore structure, whether directional cues persist over supracellular length scales sufficient to organize cells has not been demonstrated in granular systems.

We reasoned that granular scaffolds comprised of high aspect ratio fibrous building blocks could provide locally persistent but globally variable pore-scale organizational cues, potentially enabling the kind of locally complex architecture characteristic of native tissue while retaining the advantages of granular materials. To address this, we developed a bulk fragmentation approach for the rapid fabrication of uniform, monodisperse high aspect ratio hydrogel fibers without specialized equipment. By applying orientation analysis tools from the aligned scaffold literature to granular systems, we introduce a framework for quantifying orientational persistence across length scales and demonstrate that fiber geometry controls pore structure in a manner not captured by conventional pore metrics. This persistence transmits to myotube organization through local contact guidance, demonstrated unambiguously in a globally disordered scaffold where global alignment cannot confound the result, and scaffolds with greater persistence support increased myotube elongation and multinucleation. Together these results draw from two scaffold design traditions that have developed largely in parallel, identifying pore orientational persistence as a tunable architectural parameter linking granular scaffold design to engineering locally organized tissue.

## Results

### Bulk fragmentation of physically crosslinked GelMA hydrogels yields uniform microfibers

To overcome traditional granular materials’ isotropic pore structure, we developed a fiber-based granular scaffold capable of mimicking the anisotropic tissue microstructure. By adapting a previously reported bulk fragmentation method^[43]^, we fabricated gelatin methacryloyl (GelMA)^[44]^ microfibers that can be packed and annealed to form fibrous scaffolds. Warm GelMA solution was suctioned into a syringe and allowed to solidify at 4ᵒC overnight (**Fig. 1a**). The gel was then manually extruded through a nylon mesh, yielding a mass of entangled hydrogel fibers. GelMA fibers were photocrosslinked after dilution in cold PBS to stabilize the fibers while minimizing annealing.

**Figure 1.**
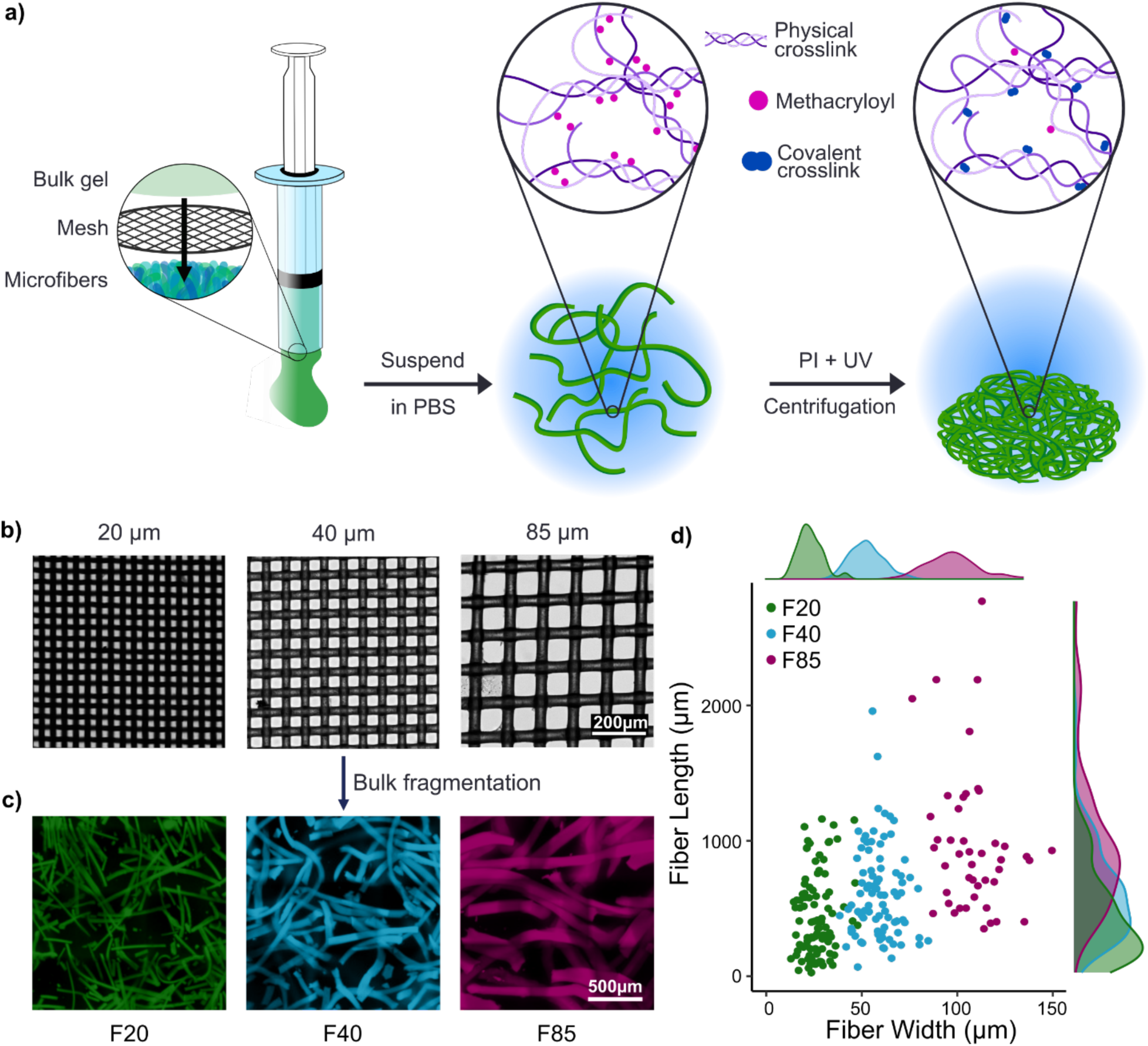
Bulk fragmentation of physically crosslinked GelMA hydrogels yields uniform microfibers. a) Schematic of the process used to fabricate and isolate GelMA microfibers. Physically-crosslinked GelMA hydrogels are extruded through a mesh, yielding individual microfibers. The microfibers are suspended in PBS and photoinitiator (PI) to create a dilute suspension prior to ultraviolet (UV) light crosslinking, reducing the likelihood of premature annealing. Covalently-crosslinked microfibers are packed via centrifugation. b) By varying the aperture size of the mesh, fibers of different widths could be fabricated as seen in panel c). d) Quantification of fiber width shows minimal overlap between fibers extruded from 20, 40, and 85 µm aperture meshes, indicating that mesh aperture size effectively controls fiber diameter. Mean fiber length also increased with aperture size, though length still varied greatly within groups.

Fiber width was easily controlled by changing the mesh aperture size (**Fig. 1b**). GelMA fibers extruded through 20 µm, 40 µm, and 85 µm meshes yielded fibers of 26.2 ± 7.1 µm (F20), 58.8 ± 9.5 µm (F40), and 108 ± 14.9 µm (F85) width respectively (**Fig. 1c-d**). Fiber length similarly increased with aperture size (426 ± 301 µm, 616 ± 336 µm, and 978 ± 531 µm for F20, F40, and F85 respectively), with fibers exceeding 1 mm in length frequently observed. These fiber dimensions were intentionally selected to be substantially larger than individual cells, with the goal of creating cell-scale porosity when fibers were packed into scaffolds. The polydispersity indices (PDI) based on fiber width shows similar narrow monodisperse distribution as microfluidic-based approaches of microparticle fabrication^[45]^, though this 1D metric may not be appropriate to adequately describe dispersity of 3D anisotropic particles (PDI of 1.07, 1.03, and 1.02 respectively). Successful fiber fabrication also depended on the open area fraction of the meshes; meshes with open area fraction of at least 25% reliably produced fibers, while lower open area fraction yielded irregular fractured particles (mesh structure comparison for 20 µm meshes seen in **Fig. S1**).

To evaluate the generalizability of this fabrication method, other hydrogel systems of similar stiffnesses were extruded through the meshes. Extrusion of covalently-crosslinked norbornene-modified hyaluronic acid yielded coarsely fractured particles (**Fig. S2a**). This was expected as bulk fragmentation of covalently-crosslinked hydrogels into irregular particles has been widely demonstrated. A hyaluronic acid hydrogel crosslinked by reversible adamantane-β-cyclodextrin complexing^[46]^ did not form fibers either, seemingly due to immediate reannealing into a disperse amorphous aggregate (**Fig. S2b**). These observations suggest that the viscoelastic and thermoreversible properties of gelatin-based hydrogels may uniquely enable mesh filaments to split the bulk gel into continuous fibers rather than irregular particles, motivating the use of GelMA fibers for all subsequent experiments. This simple fragmentation approach therefore enables rapid production of uniform, high aspect ratio hydrogel microfibers without specialized microfabrication or electrospinning equipment.

### Packed microfibers exhibit rheological properties observed in other granular systems

To establish how these fibers perform as processable granular materials, we performed rheological characterization on packed unannealed fiber constructs. The high aspect ratio of the fibers introduces the potential for extensive entanglement that could limit rearrangement and deformability relative to isotropic granular systems^[47]^. We therefore assessed whether fibers retain the shear-thinning, yield-stress, and recovery behaviors characteristic of processable granular materials^[48]^.

Steady shear measurements revealed pronounced shear-thinning behavior across all conditions (**Fig. 2a**), consistent with flow arising from rearrangement of interacting fibers under applied deformation. Stress-controlled amplitude sweeps further identified a well-defined yielding transition (**Fig. 2b**), with materials exhibiting solid-like behavior at low stress (storage modulus, G′ > loss modulus, G″) and fluid-like behavior beyond a critical stress. The corresponding yield stress values (**Fig. 2c**) confirm that these assemblies resist deformation at low applied stresses, consistent with a jammed, interaction-driven structure. To assess reversibility of these interactions, we performed cyclic deformation tests, which demonstrated substantial recovery of mechanical properties following high strain (**Fig. 2d-e**). This recovery indicates that inter-fiber interactions disrupted during flow can reform upon removal of stress, enabling the material to regain its solid-like character without permanent structural damage.

**Figure 2.**
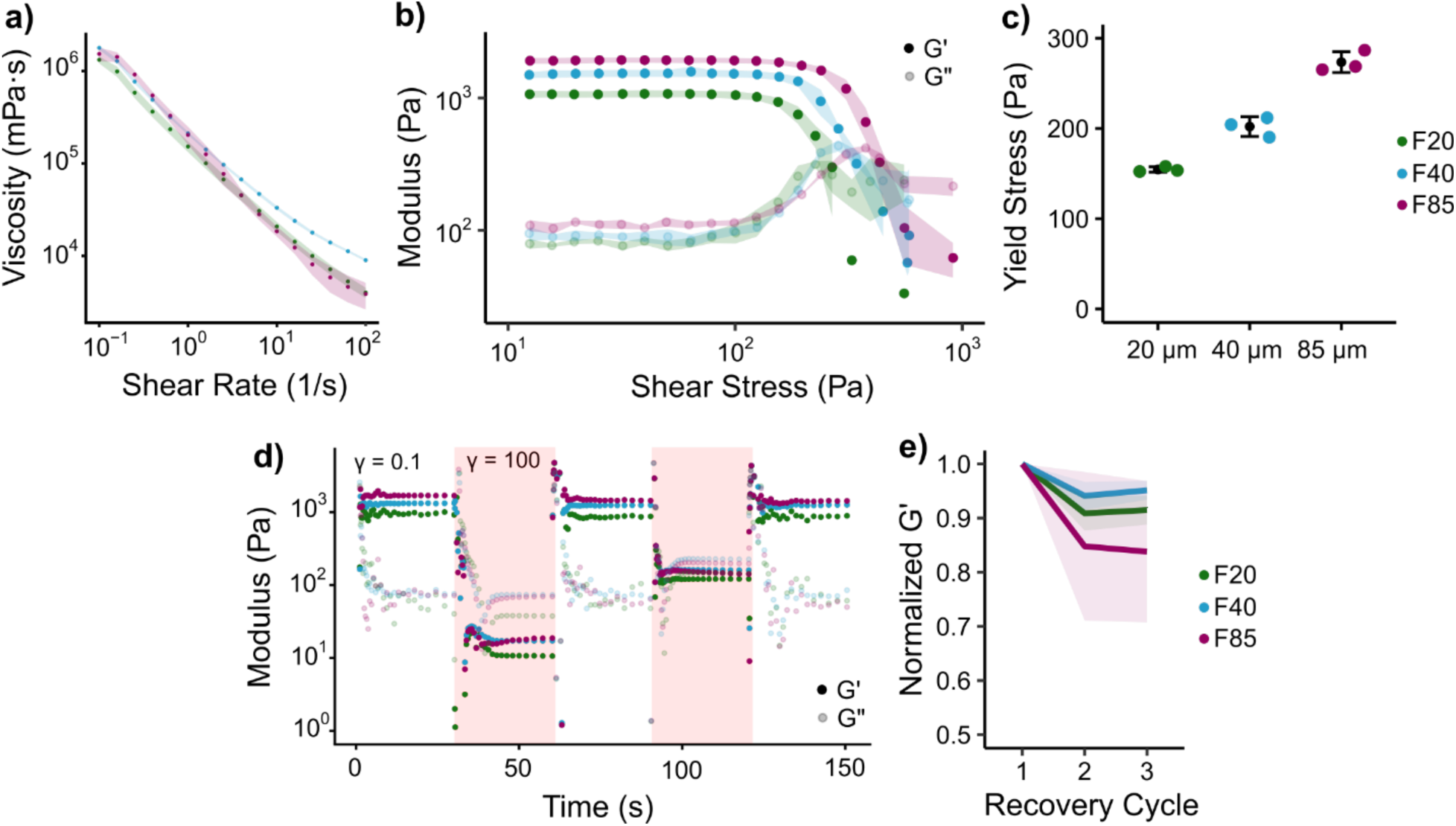
Packed fiber scaffolds exhibit granular material rheological behavior regardless of fiber width. a) Steady shear measurements show pronounced shear-thinning across all fiber widths, consistent with stress-induced fiber rearrangement. b) Stress-controlled amplitude sweeps reveal a well-defined yielding transition in all conditions, with solid-like behavior (G’ > G’’) at low stress transitioning to fluid-like behavior beyond a critical yield stress. c) Yield stress values across fiber widths, spanning a similar order of magnitude despite differences in fiber geometry. d) Cyclic deformation tests demonstrate that all scaffolds recover solid-like behavior following high strain (γ = 100%), with G’ rebounding toward pre-deformation values upon return to low strain (γ = 0.1%). e) Normalized G’ across three recovery cycles shows substantial recovery in all conditions, confirming that inter-fiber interactions disrupted during flow reform upon removal of stress. Together these results establish that packed fiber scaffolds retain the shear-thinning, yield-stress, and self-healing mechanical behavior characteristic of processable granular materials despite their high aspect ratio geometry.

Together, these results demonstrate that, despite their anisotropic geometry and potential for entanglement, packed fibers exhibit the canonical rheological signatures of granular materials. The bulk mechanical response is governed by dynamic fiber interactions and rearrangement, enabling flow under applied stress while maintaining structural integrity at rest^[6]^. Although some differences in mechanical response were observed across fiber widths, we do not draw strong conclusions regarding size-dependent trends. Variability in fiber length, packing density, and entanglement capacity, particularly as bending stiffness increases with fiber diameter, introduces multiple coupled factors that complicate direct interpretation. Accordingly, these measurements establish that packed fibers behave as mechanically jammed, rearrangeable granular materials^[48]^.

### Fiber geometry governs the spatial persistence of pore orientation

Granular scaffolds composed of isotropic particles, such as spherical microgels, typically exhibit uniform pore structures that lack a preferred directional axis. In contrast, the high aspect ratio of fibers introduces geometric anisotropy that may impart directional features to the pore space. Although these assemblies lack global alignment, it remains unclear whether fiber geometry can generate persistent directional organization. We therefore characterized pore structure across fiber sizes (F20, F40, F85) and compared these to an isotropic PEG-based MAP scaffold (spherical particle diameter of 65 µm) as a model granular material to assess how particle morphology influences pore size, organization, and the spatial persistence of orientation.

Three-dimensional reconstructions of the scaffold pore space revealed that increasing fiber width systematically alters pore structure, producing pores that are visually both larger and increasingly elongated with greater directional continuity (**Fig. 3a**). We first quantified conventional pore space descriptors to determine how fiber width modulated the local pore structure. To characterize local feature dimensions, we used local thickness analysis, a technique commonly used to characterize irregular structures like trabecular bone^[27]^. Local pore thickness at each point is defined as the diameter of the largest sphere that fits within a given phase and contains that point, providing a model-independent measure of feature size that does not assume a specific geometry. Applied to the pore phase, this yielded the mean local pore thickness, L_pore_, which increased with fiber width (12.9 ± 0.04, 17.4 ± 0.7, and 22.0 ± 2.6 µm for F20, F40, and F85 scaffolds respectively; **Fig. 3b**). Applied to the scaffold phase, local thickness yielded the mean local scaffold thickness, L_scaffold_, which similarly increased with fiber width, indicating coarsening of the solid-phase architecture within the packed scaffold (**Fig. 3c**). Despite these increases in local feature size, bulk porosity remained approximately 26% across all fibrous scaffold groups (**Fig. 3d**), indicating that wider fibers reorganized a similar total pore volume into larger local pore spaces rather than substantially altering overall void fraction.

**Figure 3.**
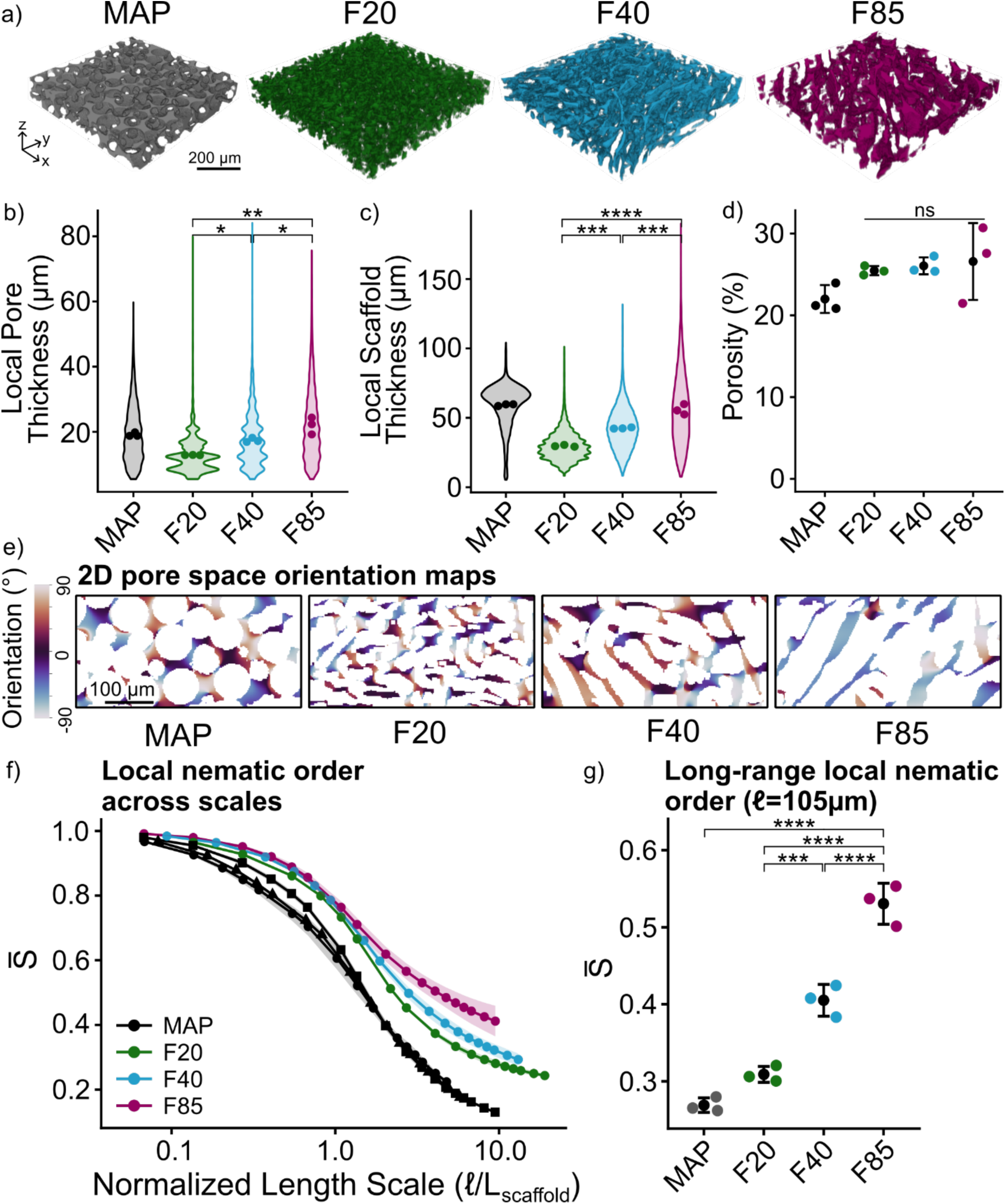
Fibers impart persistent directional organization to pore structure. a) Three-dimensional reconstructions of fiber and MAP scaffold (65 µm diameter particles) pore spaces reveal qualitative differences in pore morphology, with wider fibers producing increasingly elongated and directionally continuous pore structures (∼ 700 x 700 x 100 µm^3^ volume). b) 2D local pore thickness distributions show increasing pore size with fiber width, with MAP exhibiting comparable pore thickness to F85 despite distinct pore morphology. Violin plots depict pooled pore distributions, while points depict per-scaffold means. c) 2D local scaffold thickness (L_scaffold_) quantifies the local dimensions of the solid scaffold phase within the packed architecture. Increasing fiber width produced progressively larger local scaffold thickness, while MAP scaffolds exhibited local scaffold thickness overlapping that of F85 despite distinct scaffold morphology. d) Fibrous scaffolds exhibit similar porosity regardless of fiber width. e) Structure tensor orientation maps reveal spatially varying pore orientation in all conditions. Orientation fluctuates rapidly in MAP and thin fiber scaffolds, while wider fiber scaffolds exhibit extended regions of consistent orientation. f) Local nematic order 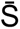 computed over increasing normalized window sizes ℓ/L_scaffold_ quantifies the spatial persistence of pore orientation. All scaffolds exhibit high local nematic order at small scales, but persistence decays more rapidly in MAP and thinner fiber scaffolds than in wider fiber scaffolds. MAP scaffolds generated from multiple particle sizes collapse onto a similar normalized persistence profile, whereas fibrous scaffolds retain elevated nematic order over larger normalized length scales, indicating that elongated fiber morphology imparts longer-range architectural persistence beyond what spherical particles can achieve. Black symbols denote MAP scaffolds of differing microstructures (square: 65 µm particles at 22% porosity; triangle: 130 µm particles at 11% porosity; circle: 130 µm particles at 15% porosity). e) Nematic order evaluated at a supracellular length scale (ℓ = 105 µm) increased with fiber width, with F85 scaffolds exhibiting significantly greater long-range orientational persistence than thinner fiber scaffolds. Despite exhibiting similar pore metrics, MAP scaffolds display substantially lower long-range nematic order than F85 scaffolds. The mean and standard deviation are shown as a single black dot with error bars. The colored dots are the per-scaffold means. Statistical analyses were performed using linear models on per-scaffold means with scaffold group as a fixed effect. Planned pairwise comparisons were evaluated using pooled variance estimates from the fitted model, analogous to Tukey’s HSD, with p-values adjusted using the Holm method.. *N* = 3 scaffolds. ****: *p* < 1×10^-4^, ***: *p* < 1×10^-3^, **: *p* < 0.01, *: *p* < 0.05.

MAP scaffolds provided an isotropic granular reference for comparison to the fibrous architectures. In contrast to the elongated pore spaces observed in wider fibrous scaffolds, MAP scaffolds exhibited more isotropic pore geometries without apparent sustained directional bias. Consistent with this particle-scale architecture, MAP local scaffold thickness was centered near the nominal 65 µm particle diameter, a correspondence further confirmed by complementary 3D local thickness analysis (**Fig. S3**). Notably, MAP and F85 exhibited similar mean pore-scale metrics, including comparable local pore and scaffold thickness, despite distinct particle geometries and pore morphologies. This suggested that differences in pore architecture may arise not only from local feature size, but also from how pores are organized across space.

Therefore, we next examined how pore orientation varies spatially using structure tensor analysis on 2D slices throughout the scaffold volume^[49]^. This approach assigns a local orientation by integrating image gradients over a defined length scale, requiring selection of an appropriate integration scale. To identify this length scale, we evaluated a range of integration lengths and found that multiple independent structural metrics converged on L_pore_ as the appropriate scale (**Fig. S4**). Accordingly, we performed structure tensor analysis at an integration scale matched to the mean 2D local pore thickness for each condition. Representative slices show that orientation fluctuates over short distances in scaffolds composed of thinner fibers, whereas larger fibers generate extended regions over which orientation remains consistent (**Fig. 3e**). MAP scaffolds exhibit high-frequency spatial heterogeneity despite comparable pore thickness to the F85 condition. These observations indicate that all scaffolds exhibit small-scale local orientational coherence but differ in how far that coherence persists across space.

To quantify the spatial persistence of orientation, we computed the local nematic order S̅, defined as the magnitude of the spatially averaged orientation field^[31,32]^, over increasing square window sizes ℓ. This metric captures how consistent orientations remain within a neighborhood defined by ℓ: high S̅ indicates that local orientations are similar, whereas low S̅ reflects spatial variability. Importantly, this metric does not quantify alignment to a single global axis, but rather the degree of local orientational consistency. A visualization of the nematic order concept can be seen in **Supplemental Video 1**, and a visualization of the calculation of orientational persistence can be seen in **Supplemental Video 2**.

When evaluated as a function of absolute window size, fibrous scaffolds exhibited fiber width-dependent persistence of pore orientation, with wider fibers retaining higher S̅ over larger real-space distances (**Fig. S5a-b**). In contrast, MAP showed the largest decrease in nematic order, consistent with the absence of sustained directional organization in spherical granular pore networks. However, because fiber width also altered local pore and scaffold dimensions, real-space persistence curves alone could not determine whether higher S̅ reflected simple architectural scaling or an intrinsic difference in how orientation was organized across the scaffold.

We therefore normalized window size by mean local scaffold thickness, L_scaffold_, to determine whether differences in persistence reflected simple rescaling of a common architecture or inherently distinct spatial organization. Normalization by local pore thickness produced similar qualitative trends (**Fig. S5c-d**), while ℓ/L_scaffold_ was used for the primary analysis because it compares persistence relative to the local feature size of the scaffold particles (**Fig. 3f and Fig. S5e-f**). After normalization, fibrous scaffolds retained distinct persistence profiles, with wider fibers maintaining elevated S̅ over greater normalized distances. Thus, fibrous architectures were not simply scaled versions of one another; increasing fiber width altered the relative length scale over which pore orientation was spatially maintained.

To determine whether the rapid decay observed for MAP reflected a particle size-specific effect or a general feature of spherical granular architectures, we also analyzed MAP scaffolds with particle diameter of 130 µm (MAP130); conventional pore metrics for MAP130 and all other scaffold groups are provided in **Fig. S3**. In real space, MAP130 retained orientational order over larger window sizes than MAP, consistent with its larger particle size and characteristic feature scale. After normalization by L_scaffold_, however, MAP and MAP130 collapsed onto a similar decay profile (**Fig. 3f**). This collapse indicates that isotropic spherical architectures are largely rescaled versions of a common pore structure. Together, these results suggest that spherical isotropic granular materials are geometrically constrained to short-range organization that primarily rescales with particle size. In contrast, fibrous granular architectures break this scaling relationship, indicating that fiber morphology changes how far directional cues extend through the pore network, rather than simply increasing pore size.

Although normalized analyses reveal whether architectures differ beyond simple feature-size scaling, cells experience scaffold structure over absolute physical distances. We therefore next evaluated S̅ at a fixed length scale ℓ = 105 µm (300 pixels at 0.35 µm/pixel), corresponding to a relevant supracellular length scale over which myotubes form (**Fig. 3g**). Unlike the structure tensor integration scale L_pore_, which captures orientation at the scale of individual pores, the nematic order window size quantifies how consistently those local orientations agree across space and is therefore intentionally large relative to L_pore_. A window size of 105 µm was selected as a conservative lower bound of reported C2C12 myotube lengths (130–520 µm)^[50]^, ensuring that nematic order is evaluated at the smallest scale relevant to multicellular organization. At this length scale, local nematic order increases monotonically with fiber width, indicating that wider fibers provide more consistent directional cues over supracellular distances. In contrast, MAP exhibited significantly lower S̅ despite having mean pore-scale metrics comparable to F85, demonstrating that similar local pore characteristics do not necessarily produce similar long-range orientational persistence. Together, these results show that fiber morphology controls not only local pore dimensions, but also the distance over which directional cues are maintained within globally disordered granular scaffolds.

### Granular pore architecture facilitates myogenesis and provides local contact guidance to myotubes

A key assumption underlying the relationship between scaffold persistence and cell organization is that cells follow local pore structure. Contact guidance by physical substrate features is well-established in globally aligned systems, but local and global directional cues are identical in these systems, making it impossible to attribute cell orientation specifically to local geometry. A granular scaffold with no preferred macroscopic orientation eliminates this confoundment, and establishing that myotubes follow local pore structure provides a direct mechanistic basis for expecting pore persistence to control the length scale of myotube organization. We therefore first confirmed that granular pore architecture is permissive for myogenesis, then directly assessed whether myotube orientation reflects local pore geometry.

To confirm that granular pore architecture is permissive for myogenesis, we compared mouse C2C12 myoblast differentiation in nanoporous GelMA hydrogels to granular F40 GelMA scaffolds. C2C12s did not form myotubes in nanoporous hydrogels whereas multinucleated myotubes readily formed in fibrous scaffolds (**Fig. 4a-c**), with morphological differences evident as early as day 1 and similar initial viability between conditions (**Fig. S6**). These results confirm that pre-existing pore architecture enables cell elongation and fusion without requiring matrix remodeling, establishing the permissive environment necessary to evaluate pore-guided myotube orientation.

**Figure 4.**
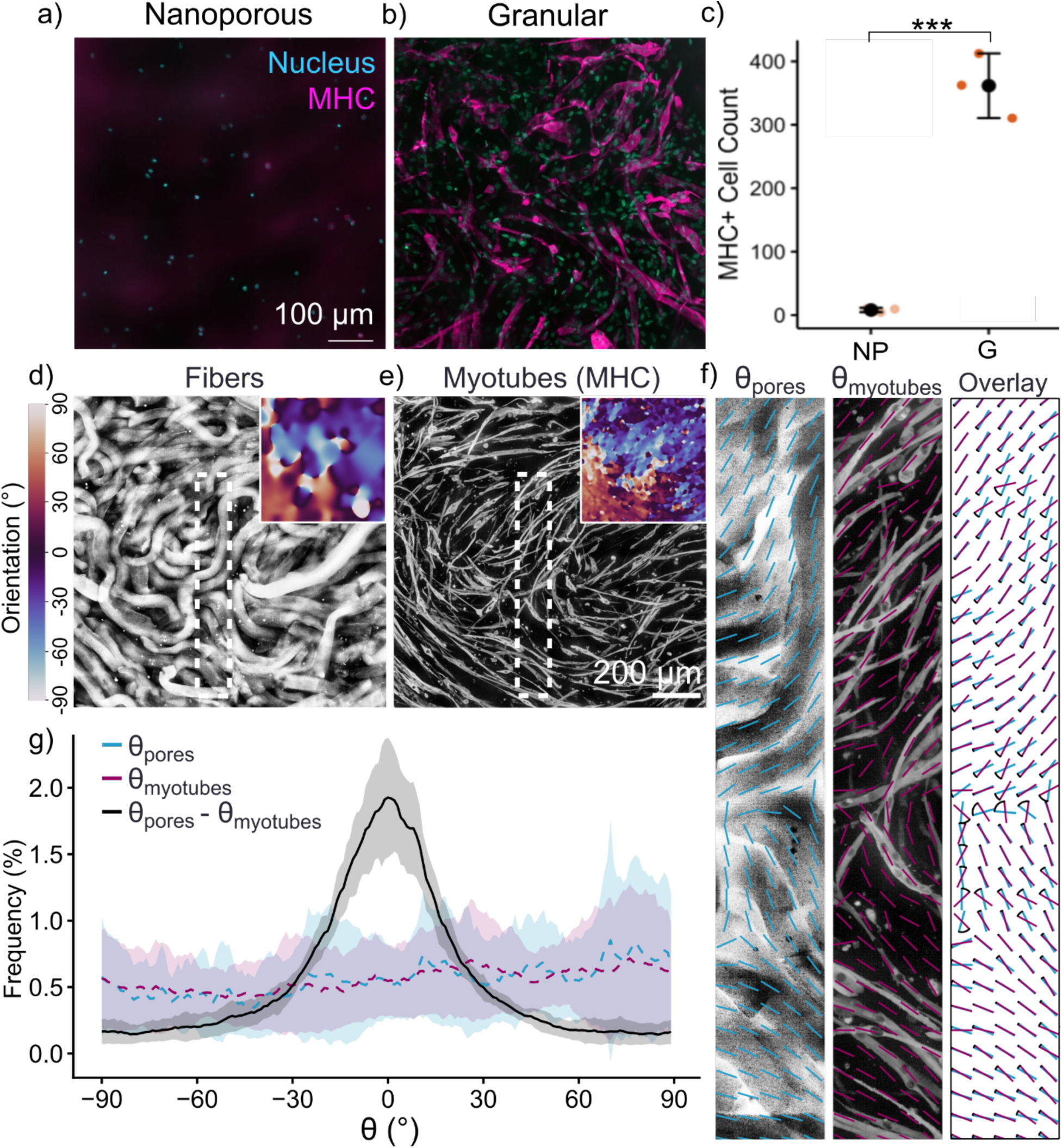
Granular pore architecture facilitates myogenesis and provides local contact guidance to myotubes. Representative confocal images of C2C12s stained for myosin heavy chain (MHC, magenta) and nuclei (cyan) after differentiation in a) nanoporous and b) granular fiber scaffolds. Myotube formation is absent in nanoporous scaffolds and robust in granular scaffolds. c) Quantification of MHC+ cell count confirms significantly greater myogenic differentiation in granular scaffolds (G) compared to nanoporous controls (NP), establishing that accessible pore architecture facilitates myogenesis in 3D. *N* = 3 hydrogels/scaffolds, ***: *p* < 1×10^-3^. Representative confocal images of d) scaffold fibers and e) MHC+ myotubes with inset orientation maps derived from structure tensor analysis. Both pore and myotube orientations are spatially variable with no preferred global axis. f) Local pore orientation vectors (θ_pores_, cyan) and myotube orientation vectors (θ_myotubes_, magenta) overlaid on the fiber image show visual correspondence between local pore and myotube directions. g) Orientation distributions of θ_pores_ and θ_myotubes_ are both uniform across angles, confirming the absence of uniaxial alignment. In contrast, the distribution of θ_pores_ − θ_myotubes_ exhibits a sharp peak at 0°, indicating that myotubes preferentially align with local pore orientation despite the lack of global scaffold order. *N* = 5 scaffolds. Together these results demonstrate that granular pore architecture directs myotube organization through local contact guidance.

To assess whether myotube orientation reflects local pore geometry, fibers and myotubes were co-stained using high molecular weight FITC-dextran and myosin heavy chain (MHC) immunostaining respectively (**Fig. 4d-e**). Consistent with disordered scaffold packing, neither fibers nor myotubes exhibited a preferred macroscopic orientation. Local orientation maps of both channels were computed using structure tensor analysis (insets of Fig. 4d-e). Orientation vectors generated by averaging these maps over 50-pixel tiles showed strong local agreement between pore and myotube orientation despite the absence of global alignment (**Fig. 4f**).

We next quantified this local organization by calculating the pixel-wise difference between fiber and myotube orientation across 15 image sets (5 scaffolds with 3 fields of view each), yielding a local misalignment metric defined as Δθ = θ_pores_ − θ_myotubes_. Because orientation is axial rather than directional, angles differing by 180° represent the same alignment axis, and misalignment was therefore calculated as the minimal angular difference between the two orientations. Although the orientation distributions of fibers and myotubes were broad due to the lack of global scaffold alignment, the distribution of Δθ was sharply concentrated around 0° (**Fig. 4g**). In most regions, pore-myotube misalignment remained within ± 30°, indicating that myotube orientation is locally constrained by the geometry of the scaffold pore structure. Correspondingly, the variance of the misalignment distribution was substantially smaller than that of either fiber or myotube orientation alone. Together, these results demonstrate that myotube organization within granular fibrous scaffolds is governed by spatially persistent structural cues, even in the absence of global alignment.

### Pore directional persistence supports increased myotube length and maturation

Having established that myotube orientation is governed by local pore structure, we next asked whether the spatial extent of myotube organization scales with scaffold persistence, the central prediction of the persistence framework. To do this, we assessed myogenic outcomes in all three granular scaffold groups with varying fiber width/pore geometry (F20, F40, and F85). Human muscle progenitor cells (hMPCs) were selected for this experiment to assess whether persistence-dependent organizational differences generalize beyond the C2C12 model to primary human cells. Myotubes formed in all conditions, with qualitative differences in morphology apparent across scaffold types. Myotubes in F85 scaffolds appeared longer and more multinucleated than those in F20 scaffolds (**Fig. 5a**). Myotube directional persistence, quantified using the same multiscale nematic framework applied to scaffold architecture, increased monotonically with fiber width. The F85 scaffolds produced myotubes with greater orientational coherence across length scales than F20 and F40 conditions (**Fig. 5b**), mirroring the scaffold architectural differences established in Figure 3.

**Figure 5.**
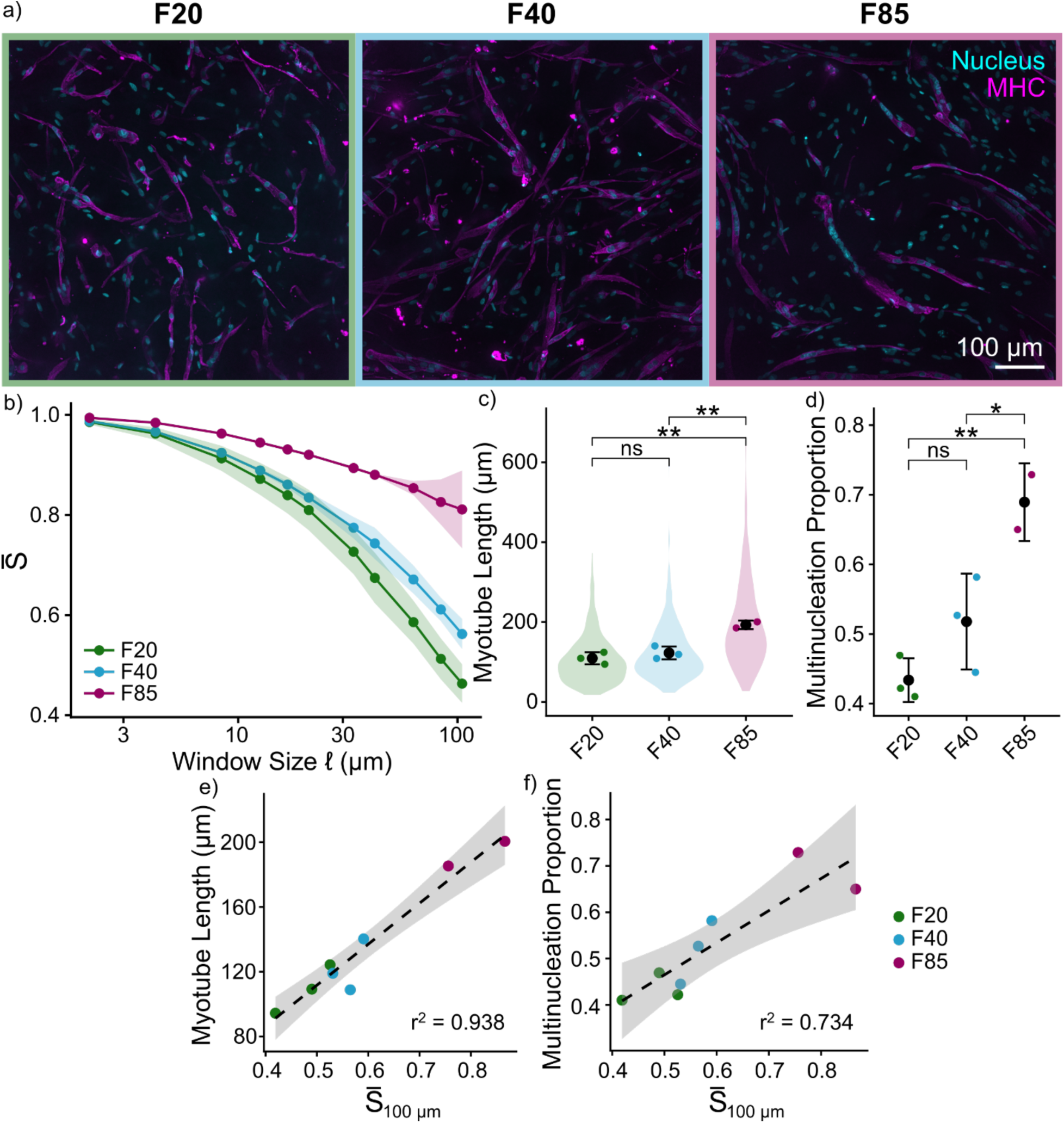
Pore orientational persistence is associated with myotube organization and maturation. a) Representative confocal images of human muscle progenitor cells stained for myosin heavy chain (MHC, magenta) and nuclei (cyan) after differentiation in F20, F40, and F85 scaffolds. Myotubes in wider fiber scaffolds appear qualitatively longer and more multinucleated. b) Myotube nematic order evaluated across length scales mirrors scaffold architectural differences, with F85 scaffolds producing myotubes with greater directional persistence than F20 and F40 conditions across the full range of measured scales. Mean is depicted by line-connected points, and standard deviation is indicated by shaded region around mean. c) Myotube length is significantly greater in F85 scaffolds compared to F20 and F40. Violin plots depict pooled length distributions, while colored points depict scaffold-wise means. d) Multinucleation proportion, defined as the fraction of MHC+ cells containing more than one nucleus, is significantly greater in F85 scaffolds than in both lower persistence conditions, indicating more advanced myogenic maturation. e-f) Myotube length and multinucleation proportion both increase continuously with myotube orientational coherence at a supracellular length scale (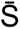_100_ μm) across all conditions, suggesting that scaffold-driven differences in spatial organization are associated with differences in myotube morphology and maturation. Linear regression and corresponding 95% confidence interval are shown as dotted lines and shaded regions respectively. The mean and standard deviation are shown as a single black dot with error bars. The colored dots are the mean metric of each scaffold. Statistical analyses were performed on the scaffold means via ANOVA followed by post-hoc Tukey’s HSD tests. *N* = 2-3 scaffolds. **: *p* < 0.01, *: *p* < 0.05.

To evaluate whether scaffold architecture and the resulting differences in myotube organization are associated with myogenic maturation, we next quantified myotube length and multinucleation across conditions. Myotube length was significantly greater in F85 scaffolds compared to both F20 and F40 conditions (**Fig. 5c**). A similar trend was observed for multinucleation, quantified as the proportion of MHC+ myotubes containing more than one nucleus. F85 scaffolds yielded a significantly higher fraction of multinucleated myotubes than both F20 and F40 conditions, while F20 and F40 did not differ significantly (**Fig. 5d**). Notably, myotube width did not differ significantly across conditions despite increasing local pore thickness with fiber width (**Fig. S7**), suggesting that myotube morphology is not simply a consequence of cells expanding to fill available cross-sectional space.

To assess whether myotube organization is associated with these morphological outcomes, myotube length and multinucleation were plotted as a function of orientational coherence at a representative supracellular length scale (100 µm). Both metrics increased with increasing coherence across conditions (**Fig. 5e-f**), indicating that more spatially coordinated myotube organization is associated with enhanced elongation and maturation. Together, these results indicate that pore orientational persistence, which governs the spatial extent of myotube organization through local contact guidance, is associated with myogenic maturation outcomes, establishing persistence as a biologically consequential architectural parameter in granular scaffold design.

## Discussion

Granular scaffolds have remarkable capacity to support tissue regeneration as demonstrated by robust cell infiltration, favorable immunomodulation, and substantial tissue formation across injury models^[1,10,11]^. However, functional recovery in anisotropic tissues has remained elusive, attributed directly to isotropic pore structure which supports cell infiltration but provides no directional guidance for coordinated multicellular organization^[10,11]^. This result reframes the design problem: the limiting variable is not regenerative capacity but the ability to provide directional cues that persist over distances sufficient to guide supracellular organization. The aligned scaffold literature has developed structure tensor analysis and nematic order metrics to quantify pore orientation, but these have had no foothold in granular scaffold design because isotropic building blocks made orientation a non-question. The result is that two fields with complementary expertise have developed largely independently, with no shared metric for comparing architectural properties across scaffold classes. Applying this analytical tradition to fibrous granular scaffolds reveals that persistence is not binary (globally aligned or absent) but a continuously tunable property that fibrous building blocks can systematically control. Fibrous granular scaffolds occupy an intermediate regime between globally aligned scaffolds, which maintain orientational coherence at all length scales, and isotropic granular materials, which lose directional organization at the cellular scale (**Fig. 6**), precisely the regime that most native tissues occupy.

**Figure 6.**
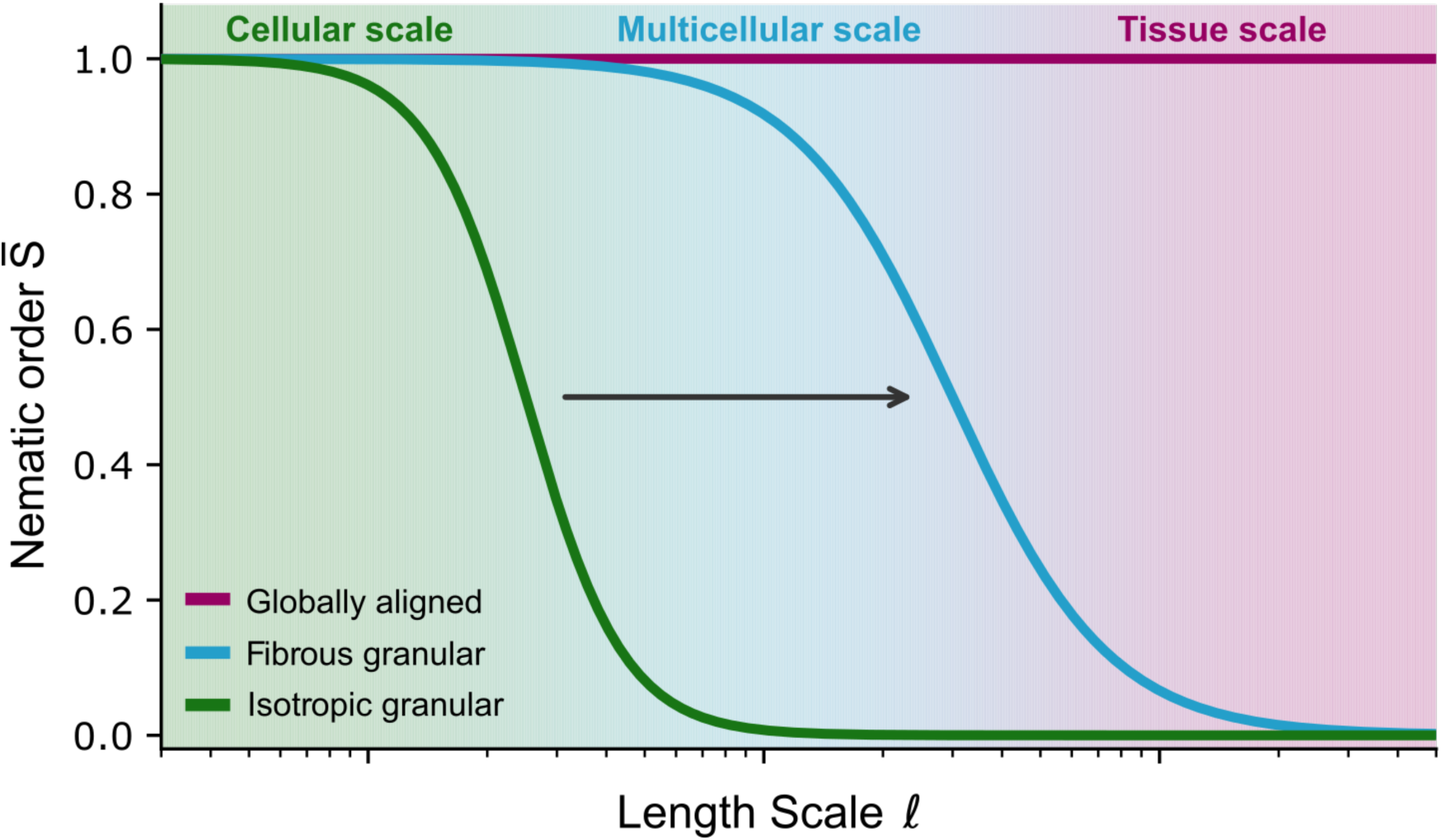
Fiber morphology extends persistence of structural cues in granular materials to the multicellular scale. Fibrous granular scaffolds occupy an intermediate architectural regime between globally aligned scaffolds, which maintain orientational coherence at all length scales, and isotropic granular materials, which lose directional organization at the cellular scale. This is precisely the domain that most native tissues occupy. The capacity of fibrous granular scaffolds to maintain orientational coherence beyond the cellular scale raises the possibility that persistence could be extended further through processing approaches, potentially enabling granular materials to achieve large-scale structural organization inaccessible through either isotropic or globally aligned scaffold designs.

Access to this intermediate regime is governed by particle morphology. Spherical building blocks produce pore geometries in which orientation varies randomly from pore to pore, with orientational coherence lost over distances comparable to a single particle diameter as demonstrated by MAP scaffolds, which exhibit near-zero nematic order at cell-relevant length scales despite pore dimensions comparable to those of wider fiber scaffolds. Prior anisotropic granular systems have characterized pore architecture primarily through aspect ratio, which describes local pore geometry in isolation and does not address whether orientation is consistent between neighboring pores. Replacing spherical particles with high aspect ratio fibers introduces directional persistence that neither spheres nor low aspect ratio rods can provide. Within fibrous scaffolds, fiber width systematically extends the length scale over which orientational coherence is maintained; wider fibers produce greater persistence at supracellular length scales despite comparable bulk porosity across conditions. Differences in bending stiffness of the fiber groups could explain this result; wider fibers resist local deformation during packing, preserving orientational continuity over longer distances than thinner fibers whose greater compliance allows more local reorganization. This relationship between fiber geometry and persistence is what places fibrous granular scaffolds in the intermediate architectural regime and suggests that persistence can be tuned by controlling mechanical properties and processing conditions rather than simply varying particle size.

This intermediate architectural regime instructs cell organization through contact guidance: myotubes align with local pore structure despite the absence of any global scaffold axis, with pore-myotube misalignment sharply concentrated near zero across measured regions despite broad and uniform orientation distributions in both channels. This extends the established principle of contact guidance beyond globally ordered substrates to a disordered granular material, suggesting that local persistence rather than global alignment is the relevant requirement for directional cell guidance. Greater persistence is also associated with enhanced myogenic maturation: myotube length increased from F20 (109 ± 15 µm) to F85 (193 ± 11 µm) and multinucleation proportion increased from F20 (0.43 ± 0.03) to F85 (0.69 ± 0.06) across the persistence range. Since pore persistence and local pore thickness co-vary across conditions, whether these differences reflect persistence specifically cannot be fully resolved. However, the invariance of myotube width despite increasing local pore thickness suggests that cells are not simply responding to having more cross-sectional space. Instead persistence, rather than pore dimensions, might be the more relevant parameter governing myotube elongation specifically.

Addressing this limitation with the ultimate goal of testing whether persistence drives functional recovery *in vivo* requires decoupling persistence from pore dimensions at the material design level. The present results reveal an inherent coupling that constrains independent parameter specification: wider fibers appear to maintain orientational continuity over longer distances, possibly due to greater bending stiffness. As a result, pore size and persistence co-vary rather than exhibit independent tunability. Attempts to increase pore size or persistence independently through packing or stiffness changes introduce bulk porosity or substrate mechanics as confounding variables. This coupling between typically independent properties reflects a fundamental difference between fibrous and isotropic granular systems: in isotropic materials, particle size alone governs pore architecture, yielding predictable and reproducible structures. In fibrous systems, pore structure is an emergent property of fiber dimensions, mechanical properties, packing efficiency, and deformation history, offering a richer but more complex architectural space. The persistence framework introduced here provides a metric to navigate that space, establishing persistence alongside conventional metrics as a useful descriptor of granular scaffold architecture, and ultimately enabling design of scaffolds that will determine whether persistence translates to functional recovery of organized tissues.

## Methods

### Materials

All materials were purchased from MilliporeSigma-Aldrich unless stated otherwise.

### Gelatin methacryloyl (GelMA) synthesis

GelMA was synthesized using a previously reported protocol^[44]^. Briefly, 100 g of gelatin B (∼ 225 g bloom, Sigma-Aldrich G9391) was dissolved in 1 L of 0.25 M carbonate buffer at pH of 9.4. Methacrylic anhydride (3.2 mL, Sigma-Aldrich 276685) was added, and the reaction solution was stirred at room temperature overnight. The reaction solution was transferred to dialysis tubing (6 kDa cutoff) and was dialyzed against RO water for 4 days with three water changes per day. The purified solution was sterile filtered, frozen, and lyophilized.

### Proton nuclear magnetic resonance

^1^H NMR experiments were performed on a Bruker Neo 400 MHz NMR Spectrometer. Gelatin and GelMA samples were dissolved in D_2_O at 20 mg mL^-1^. Spectra can be seen in **Fig. S8**.

### Microfiber fabrication

GelMA dissolved in PBS (40 mg mL^-1^) was heated to 40^ᵒ^C and loaded into a syringe. The capped syringe was placed in a 4^ᵒ^C fridge overnight to solidify. After initial gelation, the tip of the syringe was removed using a razor blade, leaving just the syringe tube without any taper that would fracture the gel. A cell strainer or mesh was then pressed against the exposed gelatin and syringe walls, and the plunger was quickly pushed to fragment the gels into fibers. Fibers were quickly placed in PBS (1 part cold PBS to 1 part room temperature PBS to 1 part fibers by volume) to separate any fibers that recoalesced. Fibers were pipetted gently until no large aggregates could be seen. Cold PBS was then quickly added in excess to solidify the individual fibers (for a total volume of 10 times greater than initial GelMA volume). LAP photoinitiator was added to yield a 1 mM solution. Fibers were then exposed to 365 nm UV light at an intensity of 5 mW cm^-2^ for 5 minutes. Uncrosslinked GelMA was removed by incubating the suspensions at 40^ᵒ^C for 30 minutes. Fiber suspensions were then centrifuged at 5000 g for 5 minutes to isolate and pack the fibers, and the supernatant was removed. Fibers underwent another centrifugation step at 21,000g for 5 minutes to remove as much interstitial liquid as possible. Cell strainers/meshes used to fabricate the different fiber sizes are as follows: Inoxia stainless steel woven wire 500 mesh for F20 group, nylon Corning 431750 40 µm cell strainer for the F40 group, and PET Pluriselect pluriStrainer 435008503 85 µm cell strainer for F85 group.

### Microfiber size quantification

Microfibers were suspended in PBS with dilute FITC-labeled 2 MDa dextran (Sigma-Aldrich FD2000S). Suspensions were then imaged using a Biotek Cytation C10 confocal fluorescent microscope (Agilent) at 20x magnification with depths over multiple hundreds of microns to capture entire fibers. Images were inverted and max z-projected prior to manual length and width measurements with ImageJ.

### Rheological characterization

All rheological measurements were performed at 25°C on an Anton Paar MCR 302 using a parallel plate (8 mm diameter) for measurements on packed microfibers. A positive displacement pipette was used to accurately load the same volume of sample. Steady shear measurements were performed by sweeping shear rate from 0.1 to 100 s⁻¹. Amplitude sweeps were performed in stress-controlled mode from 10 to 1000 Pa at 1 Hz to identify the yield stress; yield stress was calculated using the tangents method at the crossover of G′ and G″. Cyclic recovery tests were performed by alternating between low strain (γ = 0.1%, 1 Hz, 30 s) and high strain (γ = 100%, 1 Hz, 30 s) intervals for three cycles; normalized G′ recovery was calculated relative to the initial low-strain G′. All rheological data were analyzed and plotted using the R package APRheoPlotR.

### Scaffold pore structure characterization

Microfibers were suspended in concentrated LAP photoinitiator (Sigma-Aldrich 900889) in PBS to yield 1 mM concentration. Fibers were repacked via centrifugation, pipetted into silicone disc molds (8 mm diameter, 1 mm thickness), and annealed by UV exposure (365 nm, 5 mW cm^-2^, 2 minutes) to introduce secondary crosslinks between adjacent fiber surfaces. Scaffolds were incubated in with dilute FITC-labeled 2 MDa dextran (Sigma-Aldrich FD2000S) on a shaker overnight. Scaffolds were then imaged using a Biotek Cytation C10 confocal fluorescent microscope (Agilent) at 20x magnification with 105 micron depth (image resolution of 0.35 µm x 0.35 µm x 4.2 µm).

Each 2D slice of the z-stack was binarized with uneven illumination correction, Gaussian filtering, adaptive thresholding, and morphological erosion and dilation. Masks were then loaded as 3D images into ImageJ and volume rendered using the 3D Viewer plugin. Local thickness of 2D and 3D pore masks was calculated using the python package localthickness. Pore orientation was characterized by structure tensor analysis applied to distance transforms of the 2D masks throughout the entire scaffold z-stack. Transforms were first smoothed with a Gaussian kernel (σ = 1 pixel) to suppress noise, and the structure tensor was computed by Gaussian-weighted integration of gradients with σ equal to the mean 2D local pore thickness L for each condition. The dominant eigenvector of the structure tensor at each pixel defines the local pore orientation θ, reported as an angle in [−90°, 90°).

### Local nematic order quantification

Local nematic order 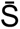 was computed as the magnitude of the spatially averaged complex orientational order parameter. For each pixel, the local orientation θ derived from structure tensor analysis was mapped to the unit complex number S = exp(2iθ), which encodes axial orientation on the unit circle. The real and imaginary components of S were each convolved with a square box filter of side length 2ℓ+1, and 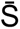 was computed as the magnitude of the resulting complex-valued average:

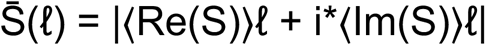

This operation averages local orientations within a square neighborhood of size (2ℓ+1)² pixels and returns a value between 0 and 1, where 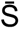 = 0 indicates isotropic orientation and 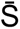 = 1 indicates perfect uniaxial alignment within the window (**Supplemental Video 1**). The box filter was applied across a range of ℓ to characterize how orientational coherence decays with increasing length scale (**Supplemental Video 2**). Window sizes were swept from 1 pixel to the full image dimension, and results were expressed both in absolute units (µm) and normalized by L (dimensionless ℓ/L). For each scaffold condition, 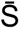 was averaged across *N* = 3 scaffolds and multiple slices per scaffold. Long-range nematic order was evaluated at a fixed window size of ℓ = 105 µm corresponding to a biologically-relevant length scale for myotube organization. Only values within the pore masks were considered when calculating 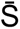.

### Cell culture

C2C12 mouse myoblasts were maintained in growth medium (DMEM supplemented with 10% FBS and 1% antibiotic-antimycotic) at 37°C, 5% CO₂. Cells were passaged at 70% confluence and used at passage numbers below 5. Differentiation was induced by switching to differentiation medium (DMEM supplemented with 2% horse serum and 1% antibiotic-antimycotic) once cells reached confluence within scaffold pores.

Human muscle progenitor cells (hMPCs, 21 year old female donor) were maintained in growth medium (DMEM/F12 1:1 supplemented with 10% FBS, 0.4 µg/mL dexamethasone, 1 µg/mL human insulin ITS-G 100x, 0.01 µg/mL hEFG, 1 ng/mL, and 0.05 mg/mL gentamycin). Differentiation was induced by switching to differentiation medium (DMEM/F12 1:1 supplemented with 10% FBS, 1 µg/mL human insulin ITS-G 100x, and 0.05 mg/mL gentamycin) once cells reached confluence within scaffold pores.

### 3D cell encapsulation

For nanoporous hydrogels, cells were resuspended in 4 wt% GelMA precursor solution at 2×10⁶ cells mL^-1^ and cast into molds. Gel precursor cell suspension was then transferred to a 4^ᵒ^C fridge for 30 minutes to physically crosslink prior to UV crosslinking (365 nm, 5 mW cm^-2^, 2 minutes), mimicking the physically and chemically crosslinked matrix of the microfibers. For granular fiber scaffolds, cell pellets were resuspended in minimal media at 50 × 10⁶ cells mL^-1^ and mixed with packed fiber pellets to yield 2×10⁶ cells mL^-1^, then transferred to molds before UV crosslinking. All constructs were cultured in growth media for 7 days prior to switching to differentiation media for 7 additional days.

### Live/Dead assay

Cell viability was assessed at day 1 and 3 post-encapsulation using a fluorescent Live/Dead kit (Invitrogen L3224) per manufacturer instructions. Briefly, constructs were incubated with calcein-AM (2 µM) and ethidium homodimer-1 (4 µM) in PBS for 30 minutes at room temperature, rinsed, and imaged immediately.

### Immunocytochemistry

Constructs were fixed in 10% formalin for 30 minutes at room temperature, and permeabilized and blocked with 0.1% Triton X-100 in 2% BSA overnight at 4^ᵒ^C. Primary antibody against myosin heavy chain (MHC; MF20 1:1000) was applied overnight at 4°C. After PBS washes, constructs were incubated with Alexa Fluor-conjugated secondary antibody (1:2000) overnight at 4°C. Nuclei were counterstained with DAPI (1 µg/mL). For fiber visualization, constructs were incubated in high MW FITC-dextran on a shaker for 2 hours. All fluorescent images were acquired on a Biotek Cytation C10 spinning disc confocal microscope using a 20x objective. Z-stacks spanning approximately 100-micron depth were acquired and maximum intensity projected for analysis.

### Myotube orientation and persistence quantification

Myotube orientation was quantified from MHC-stained maximum projection images using structure tensor analysis with integration scale matched to mean 2D local thickness of myotubes across all groups. Local nematic order of myotubes was computed using the same multiscale framework applied to pore architecture. For pore-myotube misalignment analysis, pore orientation maps were generated from FITC-dextran channel images and myotube orientation maps from MHC channel images from co-stained specimens. Pixel-wise angular difference was computed as the minimal axial difference: Δθ = min(|θ_pores_ − θ_myotubes_|, 180° − |θ_pores_ − θ_myotubes_|). Distributions of Δθ were computed across 15 image sets (5 gels × 3 fields of view per gel).

### Myotube morphology quantification

Nuclei and myotubes were segmented using the Segment Anything Model^46^ plugin SAMJ in ImageJ on maximum projected DAPI and MHC images respectively. Myotube morphological metrics including length were quantified using scikitimage function region_props() in python. Multinucleation was quantified as the proportion of MHC-positive cells containing more than one DAPI-positive nucleus, where a nucleus needed at least 70% of its area overlapping a myotube.

### Statistical analysis

All statistical analyses were performed in R. Group comparisons were made using one-way ANOVA with Tukey’s honest significant difference post-hoc test. Data are reported as mean ± standard deviation unless otherwise noted. Sample sizes are reported in figure captions.

## Acknowledgments

We thank Prof. Chris Highley for helpful discussions. We thank Prof. George Christ for his generous gift of human-derived muscle progenitor cells. MAP scaffolds were kindly provided by Prof. Don Griffin and Annabelle Hendrickson. This work was supported by the NIH (R01AR078866, R35GM162024). The content is solely the responsibility of the authors and does not necessarily represent the official views of the National Institutes of Health.

