## Supplementary Figures for "Emergent directional persistence in fibrous granular scaffolds guides myotube organization"

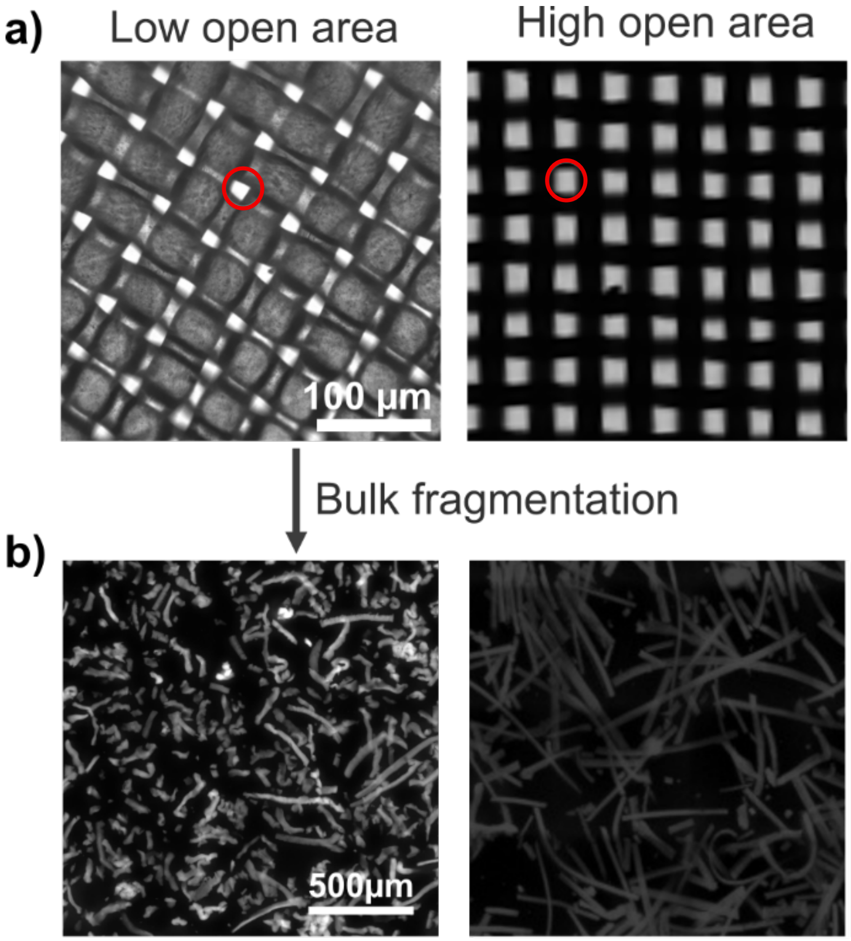


**Figure S1. Greater mesh open area fraction enables fabrication of uniform fibers without fracture.** a) Images of meshes with same aperture size as reported by the respective manufacturers. Red circles indicate location of apertures. b) Microgels fabricated from these different meshes show that low open area fraction meshes do not produce long uniform fibers. The images depicting the high open area mesh and corresponding fibers are replicated from Figure 1 (F20 fiber group).

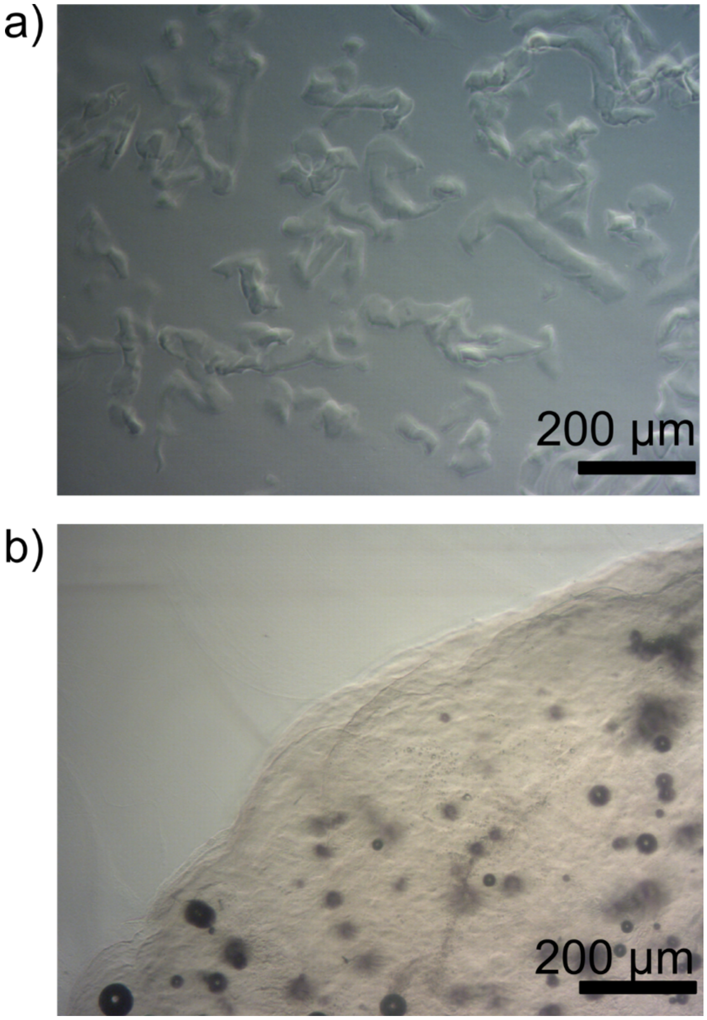


**Figure S2. Crosslinking chemistry dictates fiber fabrication potential.** a) Bulk fragmentation of a covalently-crosslinked norbornene-modified hyaluronic acid hydrogel yields highly fragmented irregular particles. b) Bulk fragmentation of a supramolecular adamantane-β-cyclodextrin crosslinked hydrogel yielded a recoalesced bulk construct.

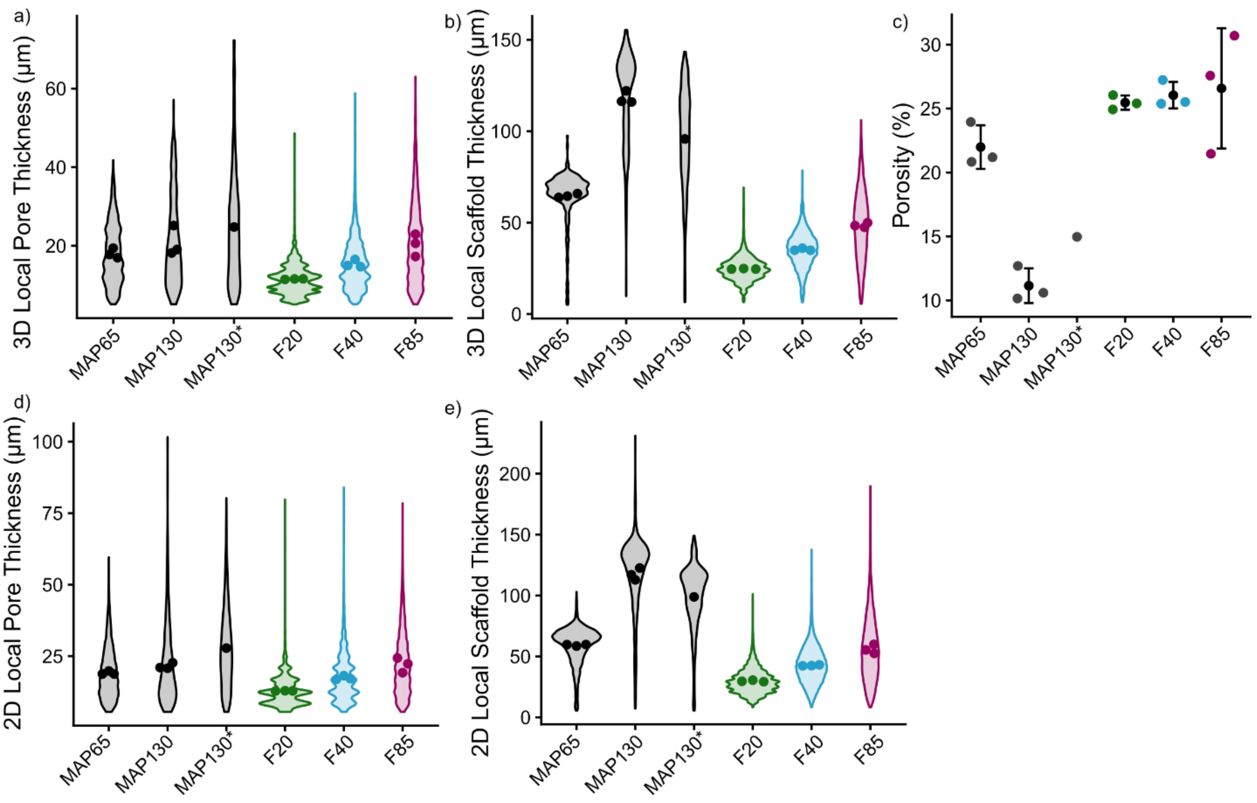


**Figure S3. Pore space and scaffold metrics across fiber scaffold groups.** Pore space and scaffold metrics shown include a) 3D local pore and b) scaffold thickness, c) porosity, and d) 2D local pore and e) scaffold thickness. Group MAP65 is the same group as MAP in Figure 3, composed of 65 μm diameter particles with porosity of ~ 22% (*N* = 3). Group MAP130 is another group of MAP scaffolds composed of 130 μm diameter particles of ~ 11% porosity (*N* = 3). Group MAP130* is a single scaffold composed of a different batch of 130 μm diameter particles with porosity of ~ 15%.

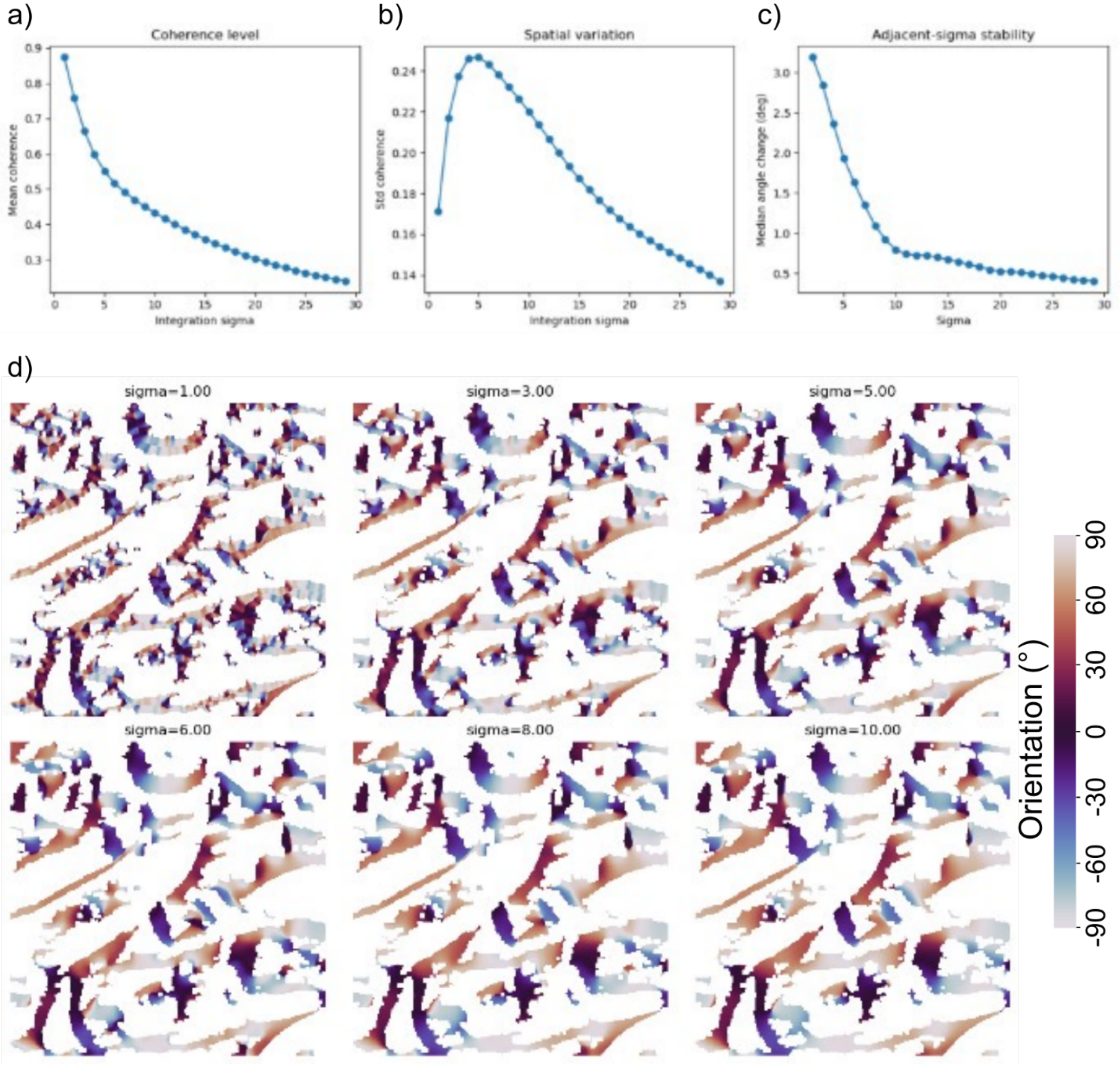


**Figure S4. Example analysis showing that L is an appropriate length scale to evaluate pore structure.** To identify appropriate characteristic length scales to evaluate structure, metric curves as a function of integration sigma were analyzed to identify elbows or peaks, indicating approximate range of possible characteristic lengths. a) Structure tensor analysis-derived coherence metric over the entire orientation map. Calculated from the eigenvalues of the pixel-wise orientations, this measure is more of a certainty metric of the orientation at each pixel. This metric is not related to the nematic order metric used in the main figure. b) Spatial variation of coherence. Structure is not adequately captured at sigmas much lower or higher than characteristic length scale, yielding low variance. As sigma approaches characteristic length scale, structure emerges and variance increases. c) Change in pixel-wise orientation as sigma increases. d) Example orientation maps at various integration sigmas. Small-scale variation decreases as sigma increases as structure is averaged over greater scales. Mean local thickness of this slice is 7.2 pixels, near the elbows of a) and c) and the peak of b).

Visually, there is minimal difference from 5-8 pixel integration sigma, indicating that local thickness is an appropriate choice for a characteristic length scale.

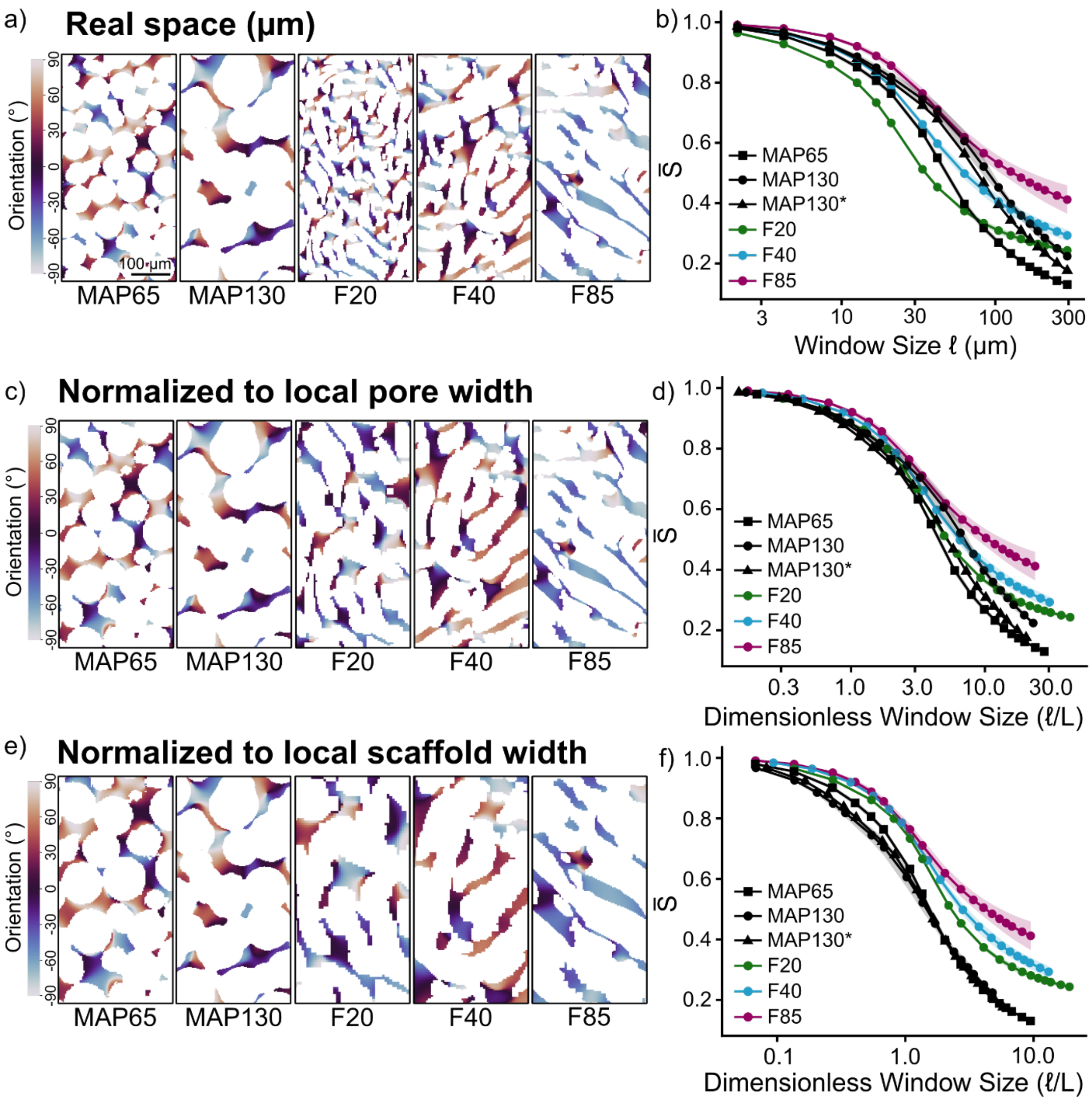


**Figure S5. Pore orientational persistence across multiple scale normalization approaches.** a,c,e) Representative pore orientation maps for MAP, F20, F40, and F85 scaffolds with images rescaled to equalize apparent scale across conditions. Nematic order curves in panel f are reproduced from Figure 3. Orientation maps in panels a, c, and e are reproduced from Figure 3 with images digitally rescaled. a) Images shown at true scale, with a 100 μm scale bar for reference. c) Images scaled by fiber width or particle diameter to theoretically equalize the building block scale. Qualitatively, neither pore thickness nor particle size appear equivalent, suggesting that this dimensionless analysis may not yield interpretable comparisons of structural differences. e) Images scaled by mean local pore thickness, the characteristic length scale of the pore structure. Both particle size and pore size appear roughly equivalent between groups, suggesting that this normalization method effectively puts scaffolds on the same scale. b,d,f) Corresponding local nematic order curves quantifying orientational persistence across window sizes under each normalization. b) Real space window size in μm reflects the absolute spatial environment experienced by cells. d) Although it likely provides limited insight into scaleless architectural differences, window size normalized to fiber width captures particle-to-particle orientational memory. For instance, the amount of structure maintained from one particle to the next is estimated at $\ell/L = 1$. f) Window size normalized to mean local pore thickness provides a scale-independent characterization of pore architecture. Across all three normalization approaches, fiber scaffolds exhibit greater orientational persistence than MAP, and F85 retains higher long-range persistence than thinner fiber scaffolds when normalized to pore width, demonstrating that persistence differences reflect genuine architectural properties rather than artifacts of scale. Similar performance of F40 and MAP groups in real space indicates that large MAP particles can impart order, but only over the scale of a single particle.

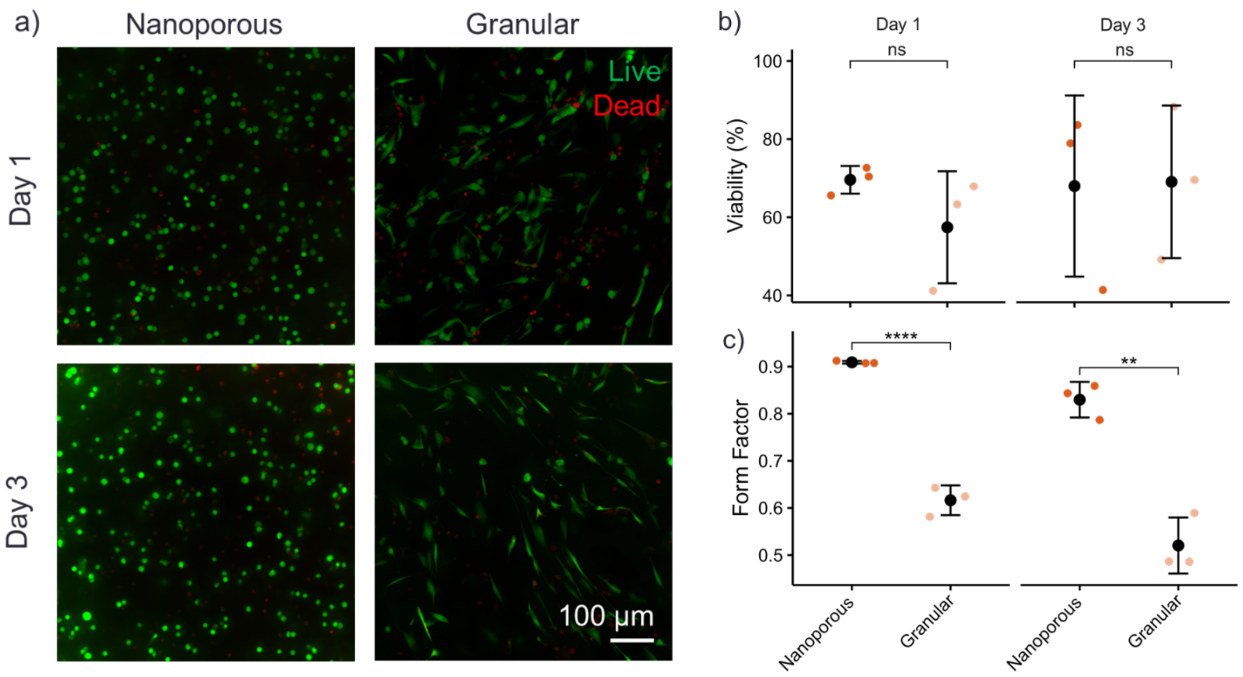


**Figure S6. Viability and cell elongation in granular fiber scaffolds.** a) Live/Dead staining of C2C12s in a nanoporous GelMA hydrogel and F40 granular GelMA scaffold. b) Viability was similar across conditions at both time points. c) Form factor, a circularity metric, shows significant cellular elongation in the granular scaffold condition even at day 1 post-encapsulation. The mean and standard deviation are shown as a single black dot with error bars. The colored dots are the mean metric of each hydrogel. Statistical analyses were performed on the hydrogel means via student's t-test. $N = 3$ hydrogels. ****: $p < 1\times10^{-5}$, **: $p < 0.01$.

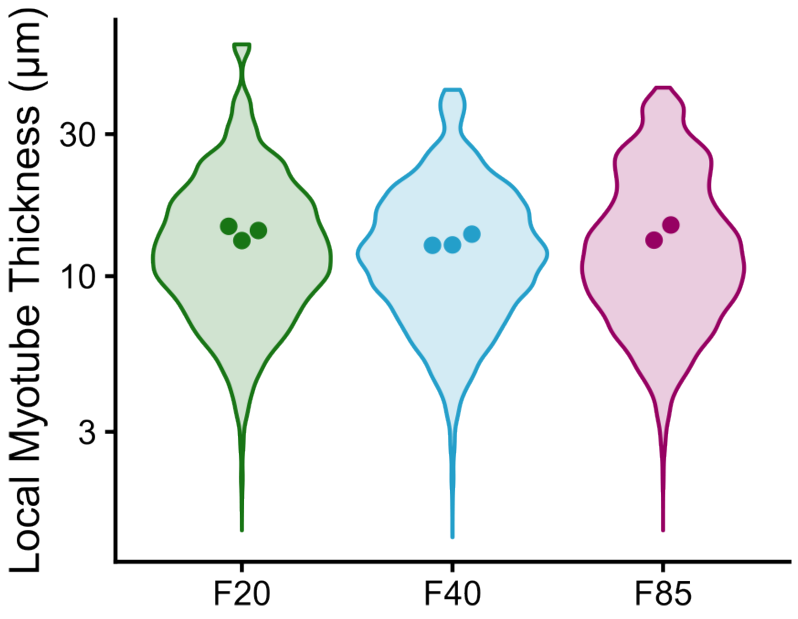


**Figure S7. Myotube width is similar in all scaffold types (ANOVA p = 0.37).**

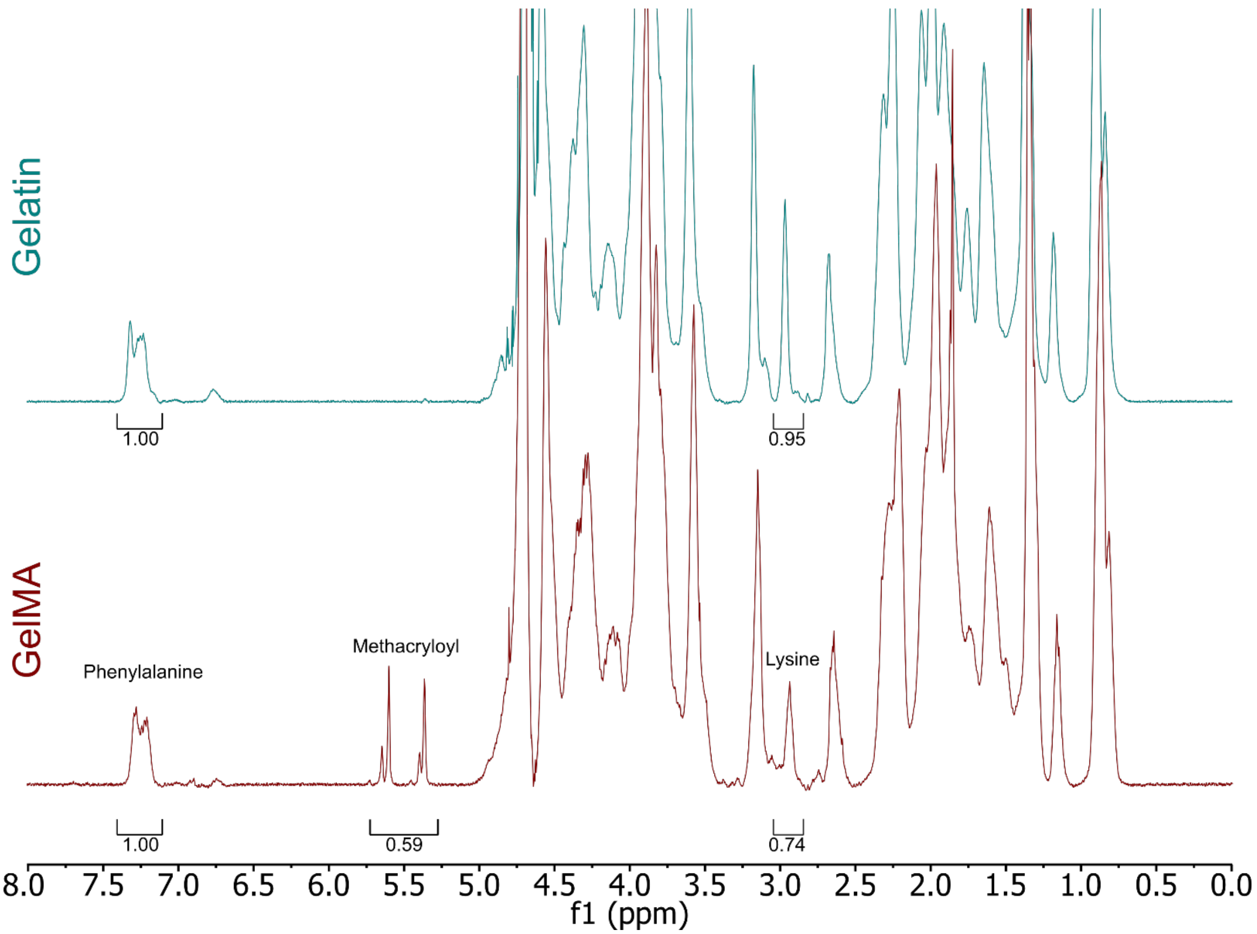


**Figure S8. $^1$H NMR spectra of gelatin and methacryloyl-modified gelatin (GelMA).** Substitution of primary amines was not complete as indicated by presence of unmodified lysine peak after modification.
